# Soil redox drives virus-host community dynamics and plant biomass degradation in tropical rainforest soils

**DOI:** 10.1101/2024.09.13.612973

**Authors:** Gareth Trubl, Ikaia Leleiwi, Ashley Campbell, Jeffrey A. Kimbrel, Amrita Bhattacharyya, Robert Riley, Rex R. Malmstrom, Steven J. Blazewicz, Jennifer Pett-Ridge

**Affiliations:** Physical and Life Sciences Directorate, Lawrence Livermore National Laboratory, Livermore, CA, 94550, USA; Department of Chemistry, University of San Francisco, San Francisco, CA, 94117, USA; DOE Joint Genome Institute, Lawrence Berkeley National Laboratory, Berkeley, CA, 94720, USA; Department of Life and Environmental Sciences, University of California Merced, Merced, CA, 95343, USA; Innovative Genomics Institute, University of California Berkeley, Berkeley, CA, 94720, USA

**Keywords:** soil viruses, soil organic matter, stable isotope probing, ^13^C, metagenomics, anoxic, oxic

## Abstract

**Background:** Wet tropical forest soils store a vast amount of organic carbon and cycle over a third of terrestrial net primary production. The microbiomes of these soils have a global impact on greenhouse gases and tolerate a remarkably dynamic redox environment—driven by high availability of reductant, high soil moisture, and fine-textured soils that limit oxygen diffusion. Yet tropical soil microbiomes, particularly virus-host interactions, remain poorly characterized, and we have little understanding of how they will shape future soil carbon cycling as high-intensity drought and precipitation events make soil redox conditions less predictable.

**Results:** To investigate the effects of shifting soil redox conditions on active viral communities and virus-microbe interactions, we conducted a 44-day redox manipulation experiment using soils from the Luquillo Experimental Forest, Puerto Rico, amended with ^13^C-enriched plant biomass. We sequenced 10 bulk metagenomes and 85 stable isotope probing targeted metagenomes generated by extracting whole community DNA, performing density fractionation, and conducting shotgun sequencing. Viral and microbial genomes were assembled resulting in 5,420 viral populations (vOTUs) and 927 medium-to-high-quality metagenome-assembled genomes across 25 bacterial phyla. Notably, over half (54%) of the vOTUs were ^13^C-enriched, highlighting their active role in microbial degradation of plant litter. These active vOTUs primarily infected bacterial phyla *Pseudomonadota*, *Acidobacteriota*, and *Actinomycetota*, and 57% were unique to a particular redox treatment. The anoxic samples exhibited the most distinct viral communities, with an increased potential for modulating host metabolism by carrying redox-specific glycoside hydrolases. However, over a third of the vOTUs were present in all redox conditions, suggesting selection for cosmopolitan viruses occurs in these soils that naturally experience dynamic redox conditions.

**Conclusions:** Our study demonstrates how redox conditions shape viral communities and virus-host interactions in soils. By applying different DNA assembly methods on stable isotope probing targeted metagenomes and incubating soils under various redox regimes, we identified distinct viral populations and observed significant variations in viral community composition and function. These findings highlight the specialized roles of viruses in microbial carbon degradation under diverse environmental conditions, providing important insights into their contributions to carbon cycling and the broader implications for climate change.

## Introduction

Tropical rainforests store about 1/3 of terrestrial organic soil carbon in 10% of global land area (Jobbagy and Jackson, 2000; Hengl et al., 2017), with some of the highest soil carbon dioxide (CO_2_) fluxes of Earth’s biomes (Raich et al. 2002). Carbon cycling in humid tropical soils is thought to be closely tied to soil moisture’s effects on microbial activity, with lower CO_2_ fluxes modeled under both drier and very wet conditions—where anoxia is presumed to limit microbial respiration (Moyano et al., 2013). However, belowground dynamics in tropical soils remain a major source of uncertainty in Earth systems models (Ciais et al. 2013; Todd-Brown et al. 2014), largely because of a lack of mechanistic data (Ghezzehei et al., 2019) on the complex relationships between soil moisture, microbial activity, and mineralogy (Cusack et al. 2023). Given the declining tropical CO_2_ sink (Brienen et al., 2015; Hubau et al., 2020), it is critical to improve our understanding of soil microbial feedbacks to climate change, and better predict how increases in temperature, episodic droughts (Janssen et al., 2020; Barkhordarian et al., 2019) and high-intensity rainfall events affect soil microbiome composition, interactions, and functionality.

Globally, soil oxygen and redox traits have a major influence on the structure and distribution of soil microbial communities (Fierer 2017; Ramírez-Flandes et al. 2019). In many humid tropical soils, the combination of ample moisture, warm conditions, frequent organic carbon inputs and fine-textured soils provide a dynamic redox landscape that oscillates between oxygenated and anaerobic conditions (Silver et al 1999; Liptzin et al., 2011), constraining both the metabolism of tropical soil microbes and interactions that regulate many aspects of soil carbon and nutrient cycling (Pett-Ridge et al. 2006; Pett-Ridge et al. 2013; Lin et al. 2018; Bhattacharyya et al., 2018; Gross et al. 2019). Indeed, in soils with frequent pulses of rainfall and organic carbon, such as those of the Luquillo Experimental Forest, Puerto Rico, high functional microbiome diversity is maintained by the highly dynamic redox environment (Pett-Ridge & Firestone, 2005; DeAngelis et al., 2010). However, it is less clear how biotic factors— such as trophic interactions—shape the interplay between environmental drivers, metabolic activity, and biogeochemical fluxes. Despite numerous studies characterizing the diverse microbial communities present in tropical soils and the influence of redox conditions on microbial metabolism (Pett-Ridge and Firestone, 2005; DeAngelis et al., 2010; DeAngelis et al., 2011; DeAngelis et al., 2013; Pett-Ridge et al., 2013; Riley et al., 2023), we have almost no information on the viruses of tropical soils, how they shape their hosts’ metabolic activity, nor how periodically low oxygen conditions affect soil virus activity.

The diversity and abundance of soil viruses are staggering, with an estimated ∼10^31^ viruses globally (Mushegian, 2020) and upwards of 10^10^ virus-like particles per gram of soil (Williamson et al. 2017; Cao et al., 2022). Soil viruses influence microbial metabolism by lysing host cells or altering host metabolism using auxiliary metabolic genes (AMG) and other forms of metabolic redirection (Zimmerman et al., 2020). While viruses in marine systems kill about 40% of bacteria daily and play a major role in global carbon cycling (Roux et al., 2016; Wilhelm and Suttle, 1999), our understanding of viral diversity, activity, host interactions, and the quantitative impact of viral lysis and metabolic redirection in soil systems remains limited (Nicolas et al., 2023; Emerson et al., 2018; Trubl et al., 2021; Carreira et al., 2024). The outcome of viral infection is to produce progeny viruses, which depends heavily on the host cell’s metabolic state; thus, viruses can be affected by the environmental redox status and the availability of terminal electron acceptors. For example, viruses isolated under anoxic conditions exhibit reduced burst size, longer latency periods, and lower infection efficiency, indicating that oxygen levels can influence viral infection dynamics (Hernández and Vives, 2020). Additionally, in oxygen minimum zones, viral communities display reduced diversity, distinct genotypes, and a preference to remain within hosts rather than exist as free virions (Cassman et al., 2012; Parvathi et al., 2018; Vik et al., 2020; Gazitúa et al., 2021). These findings provide initial signs that soil redox conditions and oxygen availability might shape viral infections and the metabolic pathways of microbial communities.

We hypothesize that changing soil redox conditions will significantly alter the composition, activity, and virus-host interactions of soil viral communities. We predict distinct viral populations and interactions will emerge under fully oxic, fully anoxic, and oscillating redox conditions, with oscillating conditions leading to increased viral activity. This heightened viral activity is expected to increase microbial death, thereby decreasing carbon degradation due to dynamic shifts in oxygen availability. To test this hypothesis, we conducted a 44-day redox manipulation experiment using soils from the Luquillo Experimental Forest, Puerto Rico. To discern between the decomposition of fresh litter and native organic matter, we supplemented the soils with ^13^C-enriched plant biomass. This amendment allowed us to conduct stable isotope probing targeted metagenomics and identify microorganisms actively using ^13^C-labeled plant litter as a growth substrate, the viruses actively infecting them, virus-host dynamics, and AMGs that could indicate increased involvement in soil carbon cycling. By contrasting static conditions—fully oxic or fully anoxic—with oscillating conditions, our treatments isolated the effects of changing soil redox regimes (oxygen availability), separate from hydrological effects of soil moisture.

## Methods

### Study Site and Soil Sampling

Our study site in the Luquillo Experimental Forest is a National Science Foundation sponsored Critical Zone Observatory and Long-Term Ecological Research site (Lat. 18°18 N; Long. 65°50′ W) with naturally dynamic soil redox regimes (Bhattacharyya et al., 2018). The soils are pH ∼5 highly weathered oxisols, and experience frequent soil oxygen depletion every 14−16 days (Silver et al., 1999; Liptzin et al., 2011). For this study, surface soils (0−10 cm) were collected from a slope location near the El Verde field station in late January 2016 (Fig. S1A). The litter layer was removed from the soil surface before soils were collected using a sterile bulk density corer and shipped overnight to Lawrence Livermore National Laboratory under ambient conditions.

On February 1, 2016, soils were gently homogenized and visible plant debris, rocks, and soil macrofauna were removed manually. Microcosms were established by adding ∼20 g of homogenized dry soil in 500 mL mason jars with modified lids for sampling and flow-through gas exchange where they were fitted with a GeoMicrobial septa (Geo Microbial Technologies, Ochelata OK) with Tygon tubing extending into the jar’s interior. To control the bulk soil redox conditions, either medical grade air (oxic) or nitrogen gas (anoxic) was delivered to the bottom of each jar at 3 mL min^−1^ and vented with a syringe needle in the septa. Soils were acclimated in the dark during a pre-incubation period of 16 days under a four-day oxic, four-day anoxic fluctuating redox regime, which in prior work was shown to best resemble natural redox oscillation conditions for these soils (Silver et al 1999; Pett-Ridge and Firestone 2005). Soils were pre-incubated to stabilize soil respiration resulting from homogenization disturbance. After the ‘pretreatment’ we harvested two microcosms which serve as a baseline Time Zero. Microcosms were then divided into four redox regime treatments: (1) static anoxic, (2) static oxic, (3) 4 days oxic/4 days anoxic (flux 4-day), (4) 8 days oxic/4 days anoxic (flux 8-day), and incubated for 44 days in the dark, with 5 replicates at the final harvest point (Fig. S1B). The microcosms were amended at T_0_ with 180 mg plant biomass that was either ^13^C labeled (97 atom%) or unlabeled ^12^C ryegrass (9 mg g soil^-1^, Isolife #U-61789, Wageningen, Netherlands), representing 7.5% of the soil’s native carbon content. Plant biomass was added to soils’ surface using a fine sifter for even dispersal across the soil surface. Sterile water was added with a glass bottle mister to achieve 50% water content (∼3.5 mL) and to aid in plant biomass incorporation in the soil. All microcosms were weighed and checked repeatedly throughout the experiment to ensure soils maintained 50% water content. Air and nitrogen (N_2_) gases were bubbled through sterile water before reaching the microcosm headspace to prevent soil drying. A handheld flow meter (Proflow 6000, Restek Corp, Bellefonte, PA) was used to check the gas efflux rate to ensure all microcosms experienced the same flow rate (3 mL min^-1^); adjustments were made as needed using upstream flow controllers (Series RM Polycarbonate Flowmeter model RMA-150-SSV, Dwyer, Michigan City, IN). During each ‘redox switch’, complete replacement of the microcosm headspace (oxic to anoxic or vice versa) took 30 minutes at an increased flow rate of 16 mL min^-1^, after this the flow was returned to the standard flow rate of 3 mL min^-1^.

Replicate microcosms from all four redox treatments were destructively harvested on day 44. Microcosm jars were opened, soils were lightly homogenized and then sampled for microbial and viral analyses. Sampling and extractions for anoxic headspace microcosms were performed within an anoxic chamber (5% H_2_ and 95% N_2_). All chemical reagents used within this chamber were prepared in advance with degassed water to preserve the Fe oxidation state.

### DNA extraction, stable isotope probing fractionation, and sequencing

DNA from each microcosm was extracted in triplicate using DNA was extracted using a modified Griffith’s protocol (Griffiths et al. 2000), then pooled for downstream sequencing. For the extraction procedure, soil (0.25 g) was added to lysing matrix E tubes (MP Biomedical, Santa Ana, CA) with 0.5 mL 50 mM EDTA/350 mM KPO_4_/0.7 M NaCl/0.5% sarkosyl (w/v), 0.5 mL phenol:chloroform:isoamyl alcohol (25:24:1, pH 8), 30 µL β-Me, 20 µL BSA (400 mg mL^-1^). Soils were bead beaten for 45 sec at 6.5 m s^-1^ in a Fast-Prep (MP Biomedicals, Santa Ana, CA). Sodium chloride (0.7 M final) and 100 µL 10% CTAB/0.7 M NaCl were added to each soil slurry before centrifugation (5 min, 16,000 x g, 4°C). After collecting the aqueous layer, soils were back extracted with NaCl (0.7 M) and 0.5 mL 50 mM EDTA/350 mM KPO_4_, vortexed and centrifuged. The combined aqueous layers were washed 1:1 with chloroform:isoamyl alcohol (24:1, Sigma Aldrich, St. Louis, MO), emulsified then centrifuged. The collected aqueous layer was mixed with 2 parts 40% PEG 8000/1.6 M NaCl and incubated on ice for 2 hours. Nucleic acids were pelted by centrifugation for 30 min (16 000 x g, 4°C). Pellets were washed with cold 70% EtOH followed by centrifugation twice. Air dried nucleic acid pellets were then resuspended in 50 µL 1x Tris-EDTA and replicate extractions were combined (150 µL total).

### SIP fractionation

DNA was density fractionated on the Lawrence Livermore National Laboratory high-throughput stable isotope probing pipeline (Nuccio et al., 2022). Ten unfractionated “bulk” samples (2 replicates from each redox treatment collected after 44 days of incubation and 2 T_0_ samples) and 86 stable isotope probing density fractions spanning control (natural abundance plant litter; 43 fractions) and ^13^C-enriched samples for each redox condition (43 fractions; Table S1). There were 2–3 replicates for each isotope and 3–6 heavy fractions (1.7315–1.742 density range per sample, excluding any fractions with < 0.4 ng/µL of DNA) from each redox condition that were submitted to the JGI for metagenomic sequencing. One ^13^C sample failed during library prep and was abandoned.

### Sequencing

Libraries were generated using either the Kapa Biosystems library preparation kit (Roche) or the Nextera XT kit (Illumina), depending on the available mass. For the Kapa Biosystems libraries, 200 ng of DNA was sheared to approximately 500 bp using a Covaris LE220 ultrasonicator. The sheared DNA fragments were size selected by double-SPRI before end-repair, A-tailing, and ligation with Illumina compatible sequencing adaptors from IDT containing a unique molecular index barcode for each sample library. For Nextera XT libraries, 2 ng of DNA was used in the tagmentation reactions, followed by 9-12 cycles of PCR. Sequencing for 95 metagenomes (Table S1) was performed on the NovaSeq (Illumina) sequencer using NovaSeq XP V1 reagent kits and an S4 flowcell, following a 2 x 151 indexed run protocol.

### Read quality control and assembly methods

The paired-end Illumina reads were trimmed and screened using BBDuk according to the documentation for BBTools (Bushnell 2014). Filtered reads were corrected using bfc (version r181) with “ bfc -1 -s 10g -k 21 -t 10, and reads with no mate pair were removed, resulting in 95 FASTQ files totaling 7.74 TB of filtered reads, representing 3.40 Tbp of sequence. The resulting reads were then assembled using two different approaches. The “single assembly” approach was performed using SPAdes assembler 3.12.0 (Bankevich et al., 2012) with the following parameters: -m 2000 --only-assembler -k 33,55,77,99,127 --meta. The “co-assembly” approach was performed with MetaHipMer (Hofmeyr et al., 2020) version 2.0.1.v2.0.0 as previously described (Riley et al., 2023).

### Recovery of viruses from metagenomes

Contigs from both assembly methods were mined for viral sequences. Viral contigs were determined with VirSorter2 v2.2.4 (--keep-original-seq --include-groups dsDNAphage, NCLDV, ssDNA, lavidaviridae --min-length 5000; Guo et al., 2021), pruned of host contamination and checked for quality and completeness using CheckV (v1.01, end-to-end default parameters; Nayfach et al., 2021), and then Virsorter2 (v2.2.4, --seqname -suffix-off --viral-gene-enrich-off -- provirus-off --prep-for-dramv --include-groups dsDNAphage, NCLDV, ssDNA, lavidaviridae --min- length 5000 --min-score 0.5) was run a second time on the combined virus and provirus sequences identified by CheckV. The resulting contigs were filtered to include only viruses ≥ 10kb. Viral contigs were also mined from the metagenomes using geNomad (v1.7.0, end-to-end, default parameters; Camargo et al., 2023). Virsorter2 and geNomad vOTUs were clustered at 95% average nucleotide identity (ANI) across 85% of the shorter contig using CheckV scripts “anicalc.py” and “aniclust.py” (--min_ani 95 --min_tcov 85 --min_qcov 0). An abundance profile of the vOTUs was estimated based on post quality-controlled read mapping at ≥ 90% ANI and covering ≥ 75% of the contig (Roux et al., 2017) using Bowtie2 (v2.4.2) (Langmead et al., 2019). Any vOTUs that did not meet the coverage thresholds were removed from the final vOTU set using filterbyname.sh from BBMap version 39.01. To focus on the differences between static redox and oscillating redox conditions, the low and high fluctuating redox conditions were combined into one dataset; hereafter referred to as “flux”. Abundance was then normalized per gigabase of metagenome and by length of the contig (Roux et al., 2017). Labeling of vOTUs by ^13^C was determined by having > 1x coverage after subtracting the ^12^C-normalized relative abundance from the ^13^C-normalized relative abundance (Trubl et al., 2021).

The final vOTU set was quality checked with QUAST (v4.4; Mikheenko et al., 2016) and CheckV (v1.01). Of the 4,520 vOTUs, 59 vOTUs were considered complete, 97 were high quality, 240 were medium quality, 3,251 were low quality, and 1,773 could not be determined (Table S2). The genes from the vOTUs were annotated with DRAM (v1.5; Shaffer et al., 2020) using both the DRAM-v (DRAM-v.py annotate –input_fasta virsorter2/for-dramv/final-viral-combined-for-dramv.fa –virsorter_afii_contigs virsorter2/for-dramv/viral-affi-contigs-for-dramv.tab, DRAM-v.py distill) and DRAM (DRAM.py annotate, DRAM.py distill) workflows to identify functional genes and AMGs. In brief, DRAM-v requires the affiliated contigs file from Virsorter2 which was obtained by running the final vOTU set through Virsorter2 (v2.2.4) with the following flags: virsorter run –seqname-suffix-off –viral-gene-enrich-off –provirus-off –prep-for-dramv –min-length 5000 –min-score 0.5. To maximize annotated viral content, we ran vOTUs from the final vOTU set that were identified using geNomad through Virsorter2 in a separate run to generate the required files for DRAM-v. For each gene on a viral contig that DRAM-v annotated as metabolic (M flag), a score, from 1 to 5 (1 being most confident), denotes the likelihood that the gene belongs to a viral genome rather than a degraded prophage region or a poorly defined viral genome boundary (Shaffer et al. 2020). DRAM-v detected 1,481 putative AMGs with a score of 1–3. To minimize the number of false positives, we considered only genes that had a gene ID or a gene description and removed genes with a ‘viral’ flag (V), transposon flag (T), viral-like peptidase (P) or attachment flag (A). AMGs that had a “F” flag (near the end of the contig) were removed if either at the very front of the scaffold (gene_position == 1) or they were the annotated gene that occurred furthest down the scaffold (gene_position == max(gene_position of all AMGs with that name)). Finally, we manually curated the AMG dataset by removing genes annotated with “ribosomal”, “peptidase”, “glycosyl transferase”, “adenylyltransferase”, “methyltransferase”, or those involved in nucleotide metabolism as previously described (ter Horst et al., 2021).

### Diversity of the vOTUs by sequencing type and redox

A collector’s curve was generated for all samples, with each redox condition highlighted, using the Vegan R package (method = “random”, 100 permutations, conditioned = TRUE, gamma = “jack2”; Oksanen et al., 2013) based on the normalized relative abundance profiles of the vOTUs. Alpha diversity metrics, including richness, Pielou’s J, and the Shannon–Wiener diversity index, were calculated using Vegan (Oksanen et al., 2013) and visualized as boxplots with ggplot2 (Wickham 2016). Each diversity metric was analyzed separately using a Kruskal-Wallis test followed by Dunn’s post hoc test (method = “bonferroni”) with the R package dunn.test (Dinno 2017). Principal coordinates analysis (PCoA) plots were created using Vegan (Oksanen et al., 2013), ggplot2 (Wickham 2016), and ape (Paradis and Schliep 2019) in R (methods: “hellinger”, “bray”, PERMANOVA with adonis2, 999 permutations, betadisper, 95% confidence ellipses, pairwise.adonis) to explore differences by sequencing type or redox condition. Bray-Curtis dissimilarity was employed to calculate distances, reflecting the high heterogeneity of vOTUs across samples, followed by a Hellinger transformation to reduce the influence of highly abundant or absent vOTUs. Heatmaps of the log-transformed, normalized relative abundances of vOTUs were generated using ggplot2 (Wickham 2016) and pheatmap (Kolde 2019) with a viridis color palette. Stacked bar plots of putative viral hosts were created using ggplot2 (Wickham 2016) and reshape2 (Wickham 2007) in R. UpSet plots were constructed with the UpSetR package (v1.4.0; Gehlenborg 2019), and Venn diagrams were made with the ggvenn package (v0.1.10; Yan 2021).

### MAG curation and virus-host linking

MAGs were reconstructed from the single assembly method by read-pairs from the biological replicates and were grouped for 8 separate co-assemblies (2 timepoints x 2 SIP fractions x 2 C isotopes) with MEGAHIT v1.1.3 (--k-min 27 --k-max 127 --k-step 10) (Li et al. 2015). Contigs greater than 1 KB were separately binned with MaxBin v2.2.4 (Wu, Simmons, and Singer 2016) and MetaBAT v2.12.1 (Kang et al., 2015). Genome bins from these two binning tools were refined using the bin_refinement module of MetaWRAP v1.2.1 (-c 50 -x 10) (Uritskiy, Diruggiero, and Taylor 2018) with CheckM v1.2.2 (Parks et al., 2015), using the CPR marker set (Brown et al., 2015). The presence of ribosomal RNAs (5S, 16S, and 23S) in each MAG was assessed with barrnap v0.9 (https://github.com/tseemann/barrnap) using default settings, and tRNAs were quantified in each MAG using tRNAscan-SE v2.0.11 (tRNAscan-SE -G, Chan and Lowe 2019) Finally, MAGs were dereplicated with dRep (v2.2.3; Olm et al., 2017) and only genome bins with at least ‘medium quality’ according to MIMAG standards (Bowers et al., 2017) were retained.

MAGs were reconstructed from the co-assembly as described previously (Riley et al., 2023). Briefly, MetaBAT version 2:2.15 (Kang et al., 2019) was used to generate the bins (metabat2 -m 3000 –minS 80 –maxEdges 500 –seed 1000), completeness and contamination were assessed with CheckM version 1.2.2 (checkm lineage_wf -x fa) (Parks et al., 2015). Taxonomy was then assigned to these MAGs and those from the single assembly method using GTDB-Tk version 2.1.1 (Chaumeil et al., 2022) and GTDB database release 214 (gtdbtk classify_wf –extension fa) (Parks et al., 2022). Bins were assigned as high, medium, or low quality based on the MiMAG standards (Bowers et al., 2017) and following rRNA and tRNA assessment as described above (Table S3). MAG coverage was determined as follows: Raw reads were mapped to medium and high-quality scaffolds with Bowtie2 (v2.4.2) and resulting bam files were filtered with BBtools (v39.01) reformat.sh (minidfilter=0.95 pairedonly=t; Bushnell 2014) to include only mappings with 95% sequence identity and then were sorted by position (samtools v1.5; Li et al., 2009). Coverage was calculated with coverM (v0.6.1; Aroney et al., 2024) using the command coverm genome –proper-pairs-only –min-read-percent-identity-pair 0.95 –min-covered-fraction 0.10 -m mean. The resulting coverage table was GeTMM (Smid et al., 2018) normalized for scaffold length and sequencing effort (Table S3). Diversity metrics (richness, Shannon-Weiner’s index, and Plieu’s evenness) were calculated using the Vegan (Oksanen et al., 2013) package in R (v4.4.0). The significance between groups was determined with a Kruskal-Wallis test and then Dunn’s post hoc test (method = “bonferroni”) with the R package dunn.test (Dinno 2017).

We used three mechanisms to predict putative hosts for these vOTUs. First, iPHoP (Integrated Phage Host Prediction; Roux et al., 2023) was used to predict putative microbial hosts for the vOTUs. Briefly, iPHoP links input vOTU sequences to a host taxon based on a combination of (i) direct sequence comparison to host genomes and CRISPR spacers, (ii) overall nucleotide composition comparison to host genomes, and (iii) comparison to phages with known hosts. Here, we first used iPHoP v1.3.3 with default parameters to predict hosts for all vOTU representatives based on the default iPHoP database (Aug_2023_pub). Next, we built a custom iPHoP database by adding 927 medium and high-quality MAGs and 27 additional MAGs from the iPHoP test dataset to the default GTDB-based (GTDB-tk v2.1.1 r214) iPHoP database (using the “add_to_db” function). We ran a second host prediction on the same vOTU representatives (iPHoP v1.3.3, default parameters except for the custom host database). Host predictions were filtered using a minimum score of 75, each vOTU was connected to the host taxon (at the family rank or if the score was ≥ 90, the genus rank if available) with the highest score across both iPHoP runs (default and custom database). Our second approach involved using a gene-sharing network to cluster vOTUs into genus-level taxonomic groups using vConTACT2 v0.9.19 using prokaryotic viral refseq v85 with taxonomy approved by The International Committee on Taxonomy of Viruses (Table S4; Jang et al. 2019). Finally, vOTUs hosts were inferred from vOTU taxonomic affiliation provided by geNomad (Camargo et al., 2023).

### Availability of data and materials

The reads for the 95 metagenomes are available on JGI Gold (proposal: Microbial Carbon Transformations in Wet Tropical Soils: Effects of Redox Fluctuation https://doi.org/10.46936/10.25585/60000880) and the sample IDs are in supplementary Table S1, and under Bioproject PRJNA539145. The 5,420 vOTU sequences are available at https://zenodo.org/doi/10.5281/zenodo.11359566. The MetaHipMer co-assembly, MAGs, and annotations, along with search and analysis tools, are available in the IMG/MER Database under the taxon object ID 3300047160.

## Results

### Characterization of viruses

We compared two assembly approaches to enhance virus detection: a single assembly, where contigs were assembled separately for each metagenomic sample, and a co-assembly, where contigs were assembled simultaneously using reads compiled from all metagenomic samples. The single assembly produced 87,863 contigs ≥ 10kb in length and the co-assembly produced 498,187 were ≥ 10kb. Both contig sets were analyzed for viral sequences using VirSorter2 and geNomad with 11,843 viral contigs ≥ 10kb from the single assembly and 29,752 viral contigs ≥ 10kb from the co-assembly. The co-assembly captured approximately three times more viral contigs compared to single assembly. To further refine these findings, contigs ≥ 10kb from each dataset were clustered into nonredundant viral populations (vOTUs). This process yielded 1,448 vOTUs from the single assembly and 21,866 vOTUs from the co-assembly. The abundance profiles of these vOTUs were assessed and normalized (see methods) and while the single assembly maintained the same number of vOTUs (1,448) after analysis, the co-assembly experienced a five-fold reduction, leaving only 4,203 vOTUs with adequate coverage. This likely reflects the co-assembly’s ability to detect rarer viruses, which, due to their lower abundance, fell below the detection limits in the metagenome. Further quality assessment of the vOTUs revealed that the co-assembly produced higher-quality results: it had a higher N50 (17,358 vs. 15,520), yielded more high-quality and complete genomes (56 vs. 3), and identified four times as many vOTU genomes ≥ 100kb (22 vOTUs vs. 5 vOTUs).

To proceed with characterization of the viral communities, a non-redundant set of vOTUs was generated from both assembly methods resulting in a final vOTU dataset of 5,420 vOTUs (1,336 from single assembly and 4,084 from co-assembly). More than half (51%) of these vOTUs could be taxonomically classified, including three realms — *Duplodnaviria*, *Monodnaviria*, *Varidnaviria*, three phyla *Bamfordvirae* (19), *Heunggongvirae* (2,819), *Loebvirae* (2), and the family *Fuselloviridae* (1) (Table S5). We applied a gene-sharing network analyses to compare these vOTUs with RefSeq (v85) viral genomes, which identified 813 viral clusters that contained 2,548 (47%) of the vOTUs (Table S4). To better understand viral roles in carbon cycling, we tracked vOTU enrichment from host degradation of ^13^C-enriched plant litter (see methods), finding that over half (54%) of the vOTUs were active.

### Viral activity and community patterns across redox conditions

A comparison of recovered viral communities across the sampled conditions shows viral communities are well-represented in each treatment (Fig. S2). The plateau observed in the diversity curve suggests that our sampling effort successfully captured the majority of viral diversity detectable through a bulk metagenome approach. We proceeded to compare the viral communities in bulk metagenomes with those in the active viral communities identified from SIP-fractionated metagenomes under different redox conditions. Notably, over one-third of the vOTUs were shared between the bulk and SIP-fractionated metagenomes, and nearly half were present across all redox conditions (referred to hereafter as cosmopolitan vOTUs; Fig. 1A; Fig. S3A). Additionally, 27% of the vOTUs were unique to specific redox treatments (oxic, anoxic, or flux), with approximately 20% present only under static conditions and 10% only under oscillating conditions. A small portion (3%) was found only at the initial time point (T_0_), and 44% were observed exclusively at day 44, indicating that over half of the viral community remained throughout the 44-day incubation period. However, the subset of the viral community that was active showed greater variability across redox conditions. Specifically, 57% of the active vOTUs were unique to a particular redox treatment, with 26% found exclusively under oxic conditions, 16% under anoxic conditions, and 15% under oscillating conditions (Fig. S3B). This pattern suggests that many viruses infecting hosts involved in plant biomass degradation are dependent on specific redox conditions for infection.

**Figure 1.**
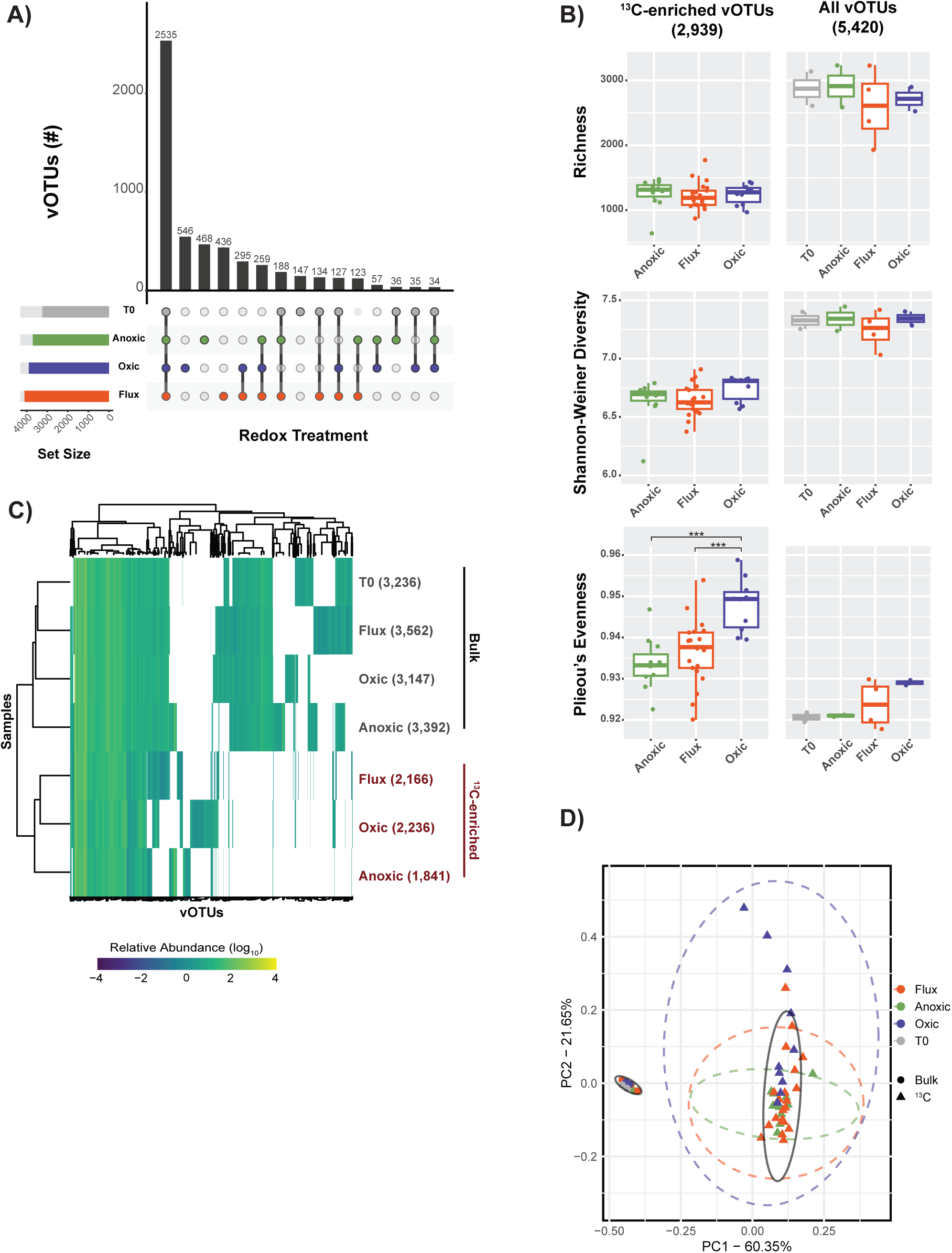
vOTU Occupancy, Diversity, and Abundance Across Redox Conditions. (A) Upset plot depicts vOTU occupancy within the different redox treatments. (B) Boxplots showing alpha diversity metrics of the ^13^C-enriched vOTUs and all vOTUs by redox condition based on vOTU normalized abundance. The median of the data are shown as solid lines in the respective boxes, the interquartile range, which spans from the first quartile (25th percentile) to the third quartile (75th percentile) is shown, and the whiskers shown the smallest and largest values within 1.5 times the interquartile ranges. (C) A heatmap showing the scaled log(10) transformed relative abundances of all vOTUs present in the bulk (top rows) samples or the ^13^C-enriched vOTUs (bottom rows). (D) Principal Coordinates Analysis of Hellinger transformation of vOTU normalized relative abundances grouped by sequencing type and redox condition. Distances were calculated with Bray-Curtis dissimilarity. The ellipses are based on 95% confidence intervals.

A comparison of viral community alpha diversity metrics between T_0_ and day 44 revealed no significant differences (Kruskal-Wallis test & Dunn’s post hoc test; p > 0.05; Fig. 1B). Similarly, comparing viral communities across different redox conditions showed no significant differences in richness or Shannon diversity, whether considering all vOTUs or just the active vOTUs (Fig. 1B). Interestingly, when assessing community evenness, oxic viral communities had higher evenness than both anoxic and T_0_ samples (though not statistically significant, Kruskal-Wallis test & Dunn’s post hoc test; p > 0.05). This trend became more pronounced in the active viral communities, where oxic samples had significantly higher evenness compared to flux and anoxic samples (Kruskal-Wallis test & Dunn’s post hoc test; p < 0.001). We further examined the relative abundance profiles of the viruses to uncover ecological patterns associated with redox conditions. The 5,420 vOTUs had coverage ranging from approximately 4x–702x, with an average coverage of 37x. Clustering the samples based on the presence and relative abundance of all vOTUs and active vOTUs for each redox condition revealed that the viral community in the flux redox condition was most similar to the T_0_ community, whereas the anoxic viral community differed the most (Fig. 1C). The T_0_ community exhibited the highest cumulative relative abundance, followed by the flux, oxic, and anoxic conditions, with the anoxic community showing approximately 26% less abundance than the T0 samples. Notably, the cosmopolitan vOTUs were the most abundant across all viral communities, regardless of the redox condition.

To further explore the differences in viral communities, we conducted an ordination analysis using vOTU Bray-Curtis dissimilarity. The resulting plot revealed significant differences between the bulk and active viral communities (PERMANOVA, p < 0.001), with no significant differences in variance within groups (betadisper ANOVA with TukeyHSD, p > 0.05; Fig. 1D) (Anderson, Ellingsen, McArdle 2006). Pairwise PERMANOVA comparisons showed no significant differences (p > 0.05) in bulk samples across redox conditions. However, significant differences were observed in the active viral communities between oxic and flux conditions (p < 0.05), oxic and anoxic conditions (p < 0.01), and anoxic and flux conditions (p < 0.01). These distinctions in viral community composition across redox conditions suggest that redox potential plays a key role in litter degradation by hosts and influences viral activity.

### Virus-host dynamics across redox conditions

To gain deeper insights into virus-host interactions under varying redox conditions, we reconstructed metagenome-assembled genomes (MAGs). Co-assembly yielded 3 high-quality and 598 medium-quality MAGs, while single-assembly produced 1 high-quality and 325 medium-quality MAGs. In total, we obtained 4 high-quality and 787 medium-quality MAGs, spanning 25 bacterial phyla and 192 genera (Table S3). The most prevalent phyla were *Pseudomonadota* (290), *Acidobacteriota* (211), *Actinomycetota* (102), *Myxococcota* (63), and *Planctomycetota* (48). A comparison of the relative abundance of these MAGs across redox gradients revealed that certain phyla thrived better under specific static redox conditions. For example, *Desulfobacterota*, *Fibrobacterota*, and *Spirochaetota* had increased relative abundances only under anoxic conditions, while *Chlamydiota* and *Bdellovibrionota* were more abundant under oxic conditions (Fig. 2A). Notably, no phyla showed higher relative abundance under fluctuating redox conditions. We further analyzed the top 25 microbial genera in each redox condition and identified 16 genera present across all redox environments, while six were abundant only under specific redox conditions (Fig. 2A).

**Figure 2.**
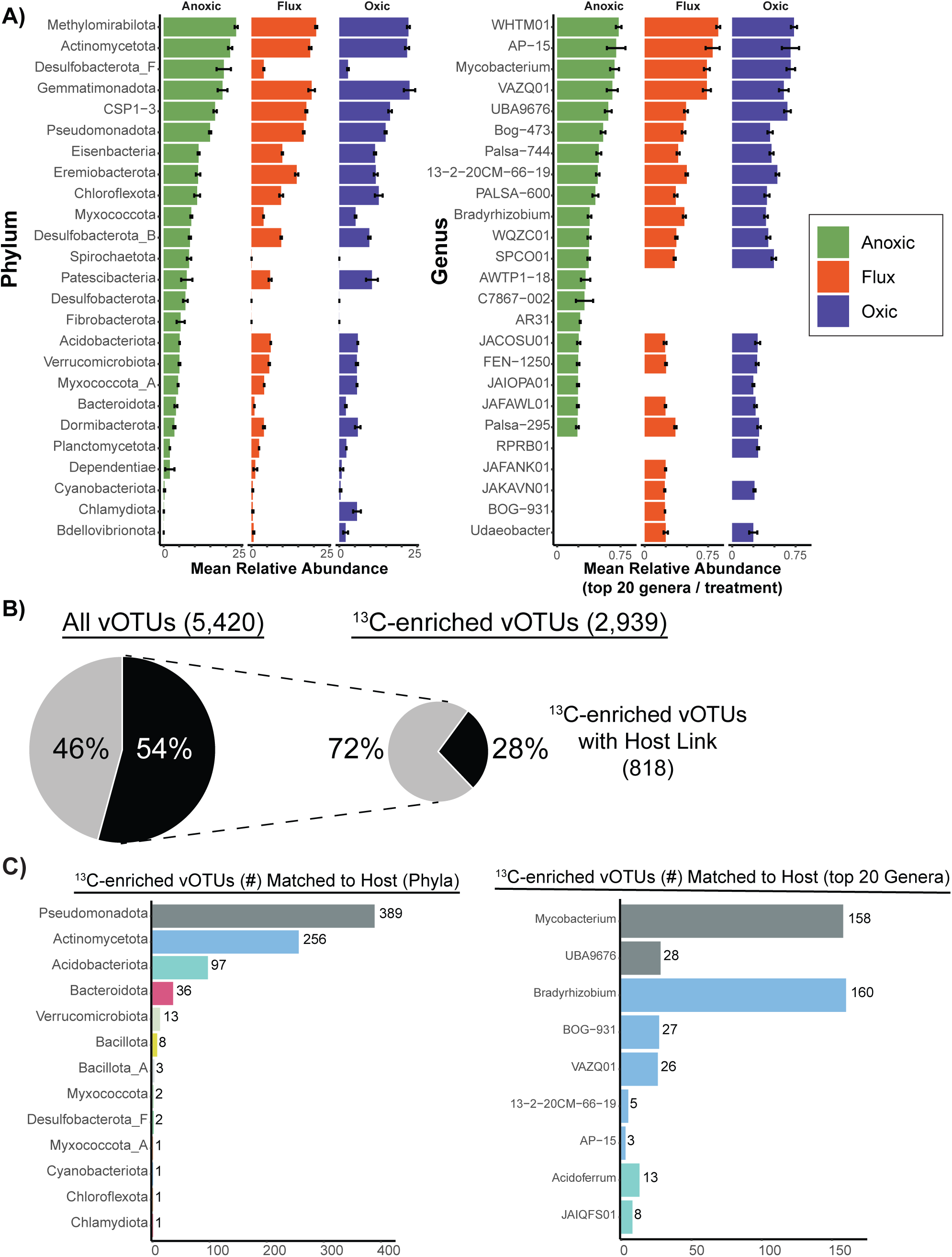
Bacterial Abundances, Virus-Host Linkages, and Taxonomic Breakdown. (A) Mean relative abundance of bacterial phyla within each treatment condition (anoxic=green, flux=orange, oxic=purple) are shown (left). Mean relative abundance of the top 20 most relatively abundant genera in each treatment condition are depicted on the right. (B) Proportion of all vOTUs (left) and all ^13^C-enriched vOTUs with a bacterial host (right). (C) Putative bacterial hosts for ^13^C-enriched vOTUs at the phylum level (left) and the genera of the top three most infected phyla (right).

To predict virus-host associations, we used the iPHoP bioinformatic tool trained on the 927 MAGs from our metagenomes, along with predictions based on viral cluster affiliations or virus taxonomy (see methods). This resulted in 13,095 virus-host linkages: 13,032 from iPHoP, 52 from VConTact2, and 11 from geNomad. Out of 5,420 vOTUs surveyed, 1,266 were linked to known hosts, with 98.3% associated with bacterial hosts (Table S6). We focused on the high-confidence virus-host linkages predicted by iPHoP, which provided host predictions for 54% of the vOTUs (Fig. 2B). A total of 538 vOTUs were linked to 341 MAGs from our data, representing 36.8% of the 927 recovered MAGs as predicted hosts. Of the total host linkages (MAGs and IMG/VR database), 3,426 had scores ≥ 90, allowing for genus-level taxonomic resolution of the predicted hosts (Table S6). Most (94%) of the putative microbial hosts for these viruses belonged to three bacterial phyla: *Actinomycetota* (43%), *Pseudomonadota* (42%), and *Acidobacteriota* (9%). Focusing on high-confidence (≥ 90 score) virus-host pairs at the genus level, we identified 111 microbial genera—five archaeal and 106 bacterial—with 53% of the putative host linkages to the genus Mycobacterium. Virus-host pairs involving active vOTUs accounted for 28% of the total (Fig. 2B), with 9,649 (74%) virus-host linkages identified overall. Of these, 818 active vOTUs had predicted hosts, including five linked to archaea (spanning five phyla and seven genera) and 803 associated with bacteria (across 13 phyla and 232 genera). Similar to the full vOTU set, bacterial hosts for the active vOTUs were predominantly from *Actinomycetota* (50%), *Pseudomonadota* (39%), and *Acidobacteriota* (8%), with most active vOTUs linked to the bacterial genera *Bradyrhizobium* and *Mycobacterium* (Fig. 2C). While we acknowledge potential database bias that may inflate *Mycobacterium* host linkages, the dominance of these two genera in the active host group may indicate an increased role in carbon turnover within these communities.

### Functional Potential of Viral Communities and Redox-Specific AMGs

To explore the functional potential of viral communities, we annotated the genomes of vOTUs (Table S7) and identified AMGs (Table S8). This analysis identified 120,315 genes, with 72.4% having unknown functions. Among the genes with known functions, 973 were predicted to be putative AMGs (score 1–3), representing 227 unique AMGs, categorized into five functional groups: 444 related to carbon utilization, 208 to energy metabolism, 140 to organic nitrogen, 65 to transporters, and 116 to miscellaneous functions, derived from 412 vOTUs (Table S8). To better understand how active viruses influence microbial metabolism under different redox conditions, we focused on AMGs from active vOTUs. We identified 139 AMGs in 143 active vOTUs, with the oxic condition having the highest number of unique active vOTUs carrying at least one AMG (n=69; Fig. 3A). However, the highest number of AMGs (n=76) were detected in vOTUs active only under anoxic conditions (Fig. 3A), indicating that viral genes with functional relevance to hosts are more prevalent under anoxic conditions. Interestingly, fewer active cosmopolitan vOTUs (n=9) were detected with AMGs (n=17) (Fig. S4A). Focusing on AMGs involved in carbon degradation, two genes related to arabinose utilization were found in two active cosmopolitan vOTUs, with higher coverage in oxic soils compared to other redox conditions. These AMGs play a role in breaking down arabinose-containing polysaccharides, contributing to the decomposition of complex organic matter (Daruwalla et al., 1981).

**Figure 3.**
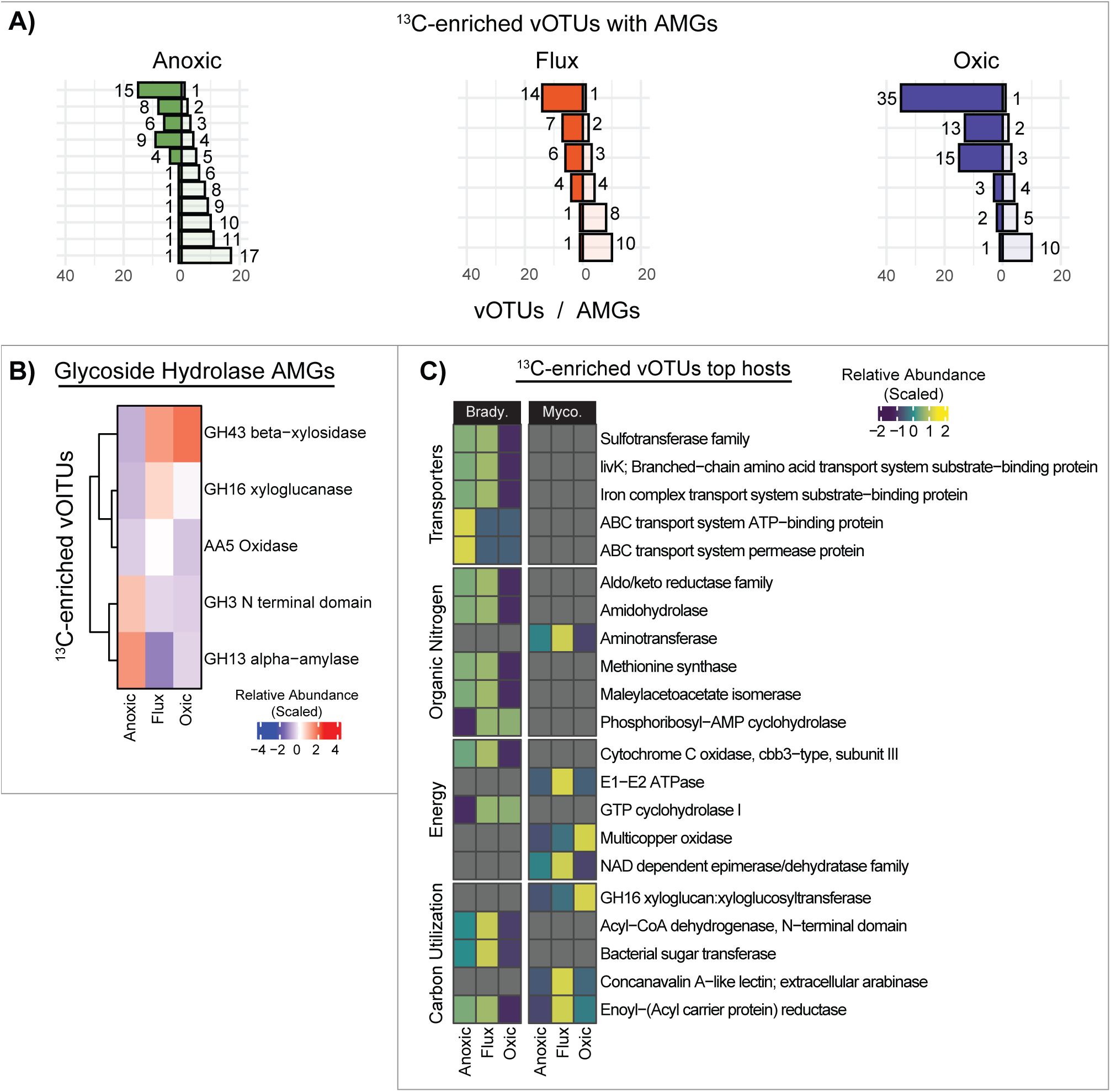
AMG characterization reveals redox specificity and vOTU roles in carbon degradation. (A) Diverging bar plots for each redox condition, where transparent bars on the right represent the number of specific AMGs detected in ^13^C-enriched vOTUs, corresponding to the solid bars on the left of each plot. The y-axis indicates the total number of distinct AMGs detected for each redox condition. (B) Heatmap showing all detected glycoside hydrolase AMGs in ^13^C-enriched vOTUs across the redox conditions. (C) Heatmap of AMGs from ^13^C-enriched vOTUs linked to genomes in the *Bradyrhizobium* or *Mycobacterium* genera.

We then examined whether non-cosmopolitan vOTUs harbored carbon degradation AMGs specific to each redox treatment. We identified 21 vOTUs containing 24 glycoside hydrolase AMGs (Fig. S4B). While no clear pattern emerged for AMGs specific to a particular redox condition, 11 active vOTUs were shown to carry 13 glycoside hydrolase AMGs. The relative abundances of active vOTUs with glycoside hydrolase families 16 and 43 genes were twice as high in oxic soils, while those with glycoside hydrolase families 3 and 13 genes were 3.2 times more abundant in anoxic soils (Fig. 3B). Further analysis of AMG content in the two most infected microbial lineages in each redox condition, *Bradyrhizobium* and *Mycobacterium*—also the most prominent hosts of active vOTUs—revealed 15 AMGs in active vOTUs infecting *Bradyrhizobium* and seven in active vOTUs infecting *Mycobacterium* (Fig. 3C). These findings highlight the potential role of viruses in enhancing the functional plasticity of their hosts across different redox environments.

## Discussion

Microbial metabolic plasticity, the ability of microbes to adjust their metabolic processes in response to varying environmental conditions, is crucial for understanding virus-host interactions. Redox conditions, influence microbial metabolic pathways, affecting their growth, survival, and ecological functions. These redox-driven metabolic shifts are essential for understanding how microbes adapt to changing environments, particularly under fluctuating oxygen levels, nutrient availability, and other geochemical gradients. The impact of redox conditions on viruses is significant because viruses rely on host metabolic processes for replication, and variations in redox states can alter viral infection dynamics, propagation, and evolution (Cassman et al. 2012; Parvathi et al. 2018; Vik et al. 2020; Gazitúa et al. 2021; Hernández and Vives, 2020). Therefore, understanding how different redox conditions affect viral community structure and virus-host interactions is critical for microbial decomposition and the fate of soil organic carbon.

Stable isotope probing targeted metagenomics is a powerful technique for tracking how specific substrates are used and identifying the microorganisms and viruses involved. Many studies use ^13^C-enriched tracers like ^13^CO_2_ or ^13^C-glucose to pinpoint substrate-specific degraders or evaluate microbial carbon use efficiency (Starr et al., 2018; Gross et al., 2019). Interestingly, viruses can also assimilate ^13^C when infecting microbes, providing insights into virus-host interactions and their role in soil carbon cycling. For example, ^13^C-CH_4_ has been used to identify viruses that target methanotrophs, which helps reduce carbon emissions (Lee et al., 2021). Another study followed ^13^C-CO_2_ through the soil food web and found ^13^C-enriched viruses, showing their role in carbon cycling in the rhizosphere (Starr et al., 2021). Recently, a study using nine different ^13^C-enriched carbon sources in soil revealed that viral activity, driven by carbon inputs, leads to microbial turnover and influences soil organic matter production (Barnett and Buckley, 2023). While these studies have advanced our understanding of carbon flow through soils and the roles of viruses, the effects of oxygen on carbon cycling and how changing redox states impact virus-host dynamics remain unclear.

### Viral and Microbial Community Dynamics Under Different Redox conditions

Assessing temporal changes in viral communities is challenging due to high spatial and physiochemical variability. Previous studies have shown that soil viral communities can exhibit seasonal variability in abundance or composition (Narr et al., 2017; Roy et al., 2020; Santos-Medellin et al., 2021; Cornell et al., 2021; Coclet et al., 2023) and undergo significant shifts over multiple years (Ter Horst et al., 2023; Sun et al., 2024). Large changes in viral community structure have also been observed following disturbances (Santos-Medellín et al., 2023; Nicolas et al., 2023). Based on these findings, we hypothesized that altering soil redox conditions—a form of disturbance—would significantly impact the composition, activity, and virus-host interactions of soil viral communities. Our results partially supported this hypothesis: we identified distinct vOTUs unique to each redox condition, indicating that some viral populations are sensitive to specific environmental changes. However, we also found a stable portion of the viral community that thrived across all redox conditions, suggesting resilience and adaptability. This aligns with previous research on microbial communities in fluctuating redox soils, which has shown that these communities are highly adapted to changing conditions and possess metabolic plasticity that allows them to persist (Pett-Ridge and Firestone, 2005). Our findings suggest that while some vOTUs are specialized for infection under specific redox conditions and show high temporal variability, cosmopolitan vOTUs maintain stability across different conditions.

Sequencing density-fractionated DNA for stable isotope probing metagenomes provides a targeted metagenomic approach that enhances the resolution of specific communities. We compared sequences from ^13^C and ^12^C metagenomes to identify vOTUs active in hosts consuming the added C substrate. We then analyzed this active vOTU pool in relation to the total detectable vOTU pool, which was determined using all sequence (bulk, ^13^C samples, and ^12^C samples) to maximize our detectable vOTU richness. We observed significant variation in viral community composition across redox conditions. Oxic samples exhibited the highest number of distinct vOTUs, while anoxic samples contained the most abundant vOTUs. Although a few ^13^C-enriched vOTUs were shared across redox conditions, the majority were unique to specific environments, suggesting that viral activity in plant biomass degradation is closely tied to redox conditions. The greater abundance of active vOTUs and AMGs in anoxic environments suggests that viruses play a more active role in carbon turnover under oxygen-limited conditions, highlighting their specialized function in modulating microbial carbon degradation across different metabolic states.

### Virus-Host Dynamics across Redox Gradients

Our study focused on how redox conditions affect viral communities and virus-host interactions leveraging and expanding on recently published MAGs from these tropical rainforest soils. Our analysis across redox gradients revealed distinct performances of certain bacterial phyla under different conditions. For instance, MAGs from phyla such as *Desulfobacterota*, *Fibrobacterota*, and *Spirochaetota* demonstrated increased relative abundances exclusively under anoxic conditions. These three phyla of bacteria include microbes that typically thrive in anoxic conditions like rumen and sediments and use anaerobic metabolism (e.g., dissimilatory sulfate reduction) to obtain energy (Diao et al., 2023; Yu et al., 2023). Other microbial populations like *Chlamydiota* (all MAGs from the order *Chlamydiales*) and *Bdellovibrionota* (all MAGs belong to the family *Bdellovibrionaceae*) thrived only under oxic conditions. Both bacterial phyla rely on other organisms for food either as obligate parasites or for predation and can contribute significantly by shaping micro-eukaryotic (e.g., amoeba) (Coulon et al., 2012) and bacterial (Hungate et al., 2021) communities. These findings align with previous work in these soils, where redox conditions can select for distinct microbial communities (Pett-Ridge and Firestone, 2005; DeAngelis et al., 2010; DeAngelis et al., 2013; Pett-Ridge et al., 2013; Riley et al., 2023).

The most recovered MAG phyla in our study were *Pseudomonadota*, *Acidobacteriota*, and *Actinomycetota*, collectively comprising 65% of the MAGs. These phyla also dominated the predicted host profiles, accounting for 95% of the virus-host linkages. Notably, the genus *Mycobacterium* (within *Actinomycetota*) was the most predicted host even though they represented less than 1% of the identified MAGs. We posit this is likely due to a combination of the fewer number of hosts predicted down to the genus level (26%) and the increased number of isolated phage-host pairs in the GTDB database (as they show promise as therapeutic agents) (Hatfull 2023). Their abundance across all redox conditions can be attributed to their diverse metabolic capabilities and functions and their metabolic plasticity, allowing them to thrive under varying environmental conditions, including different levels of oxygen availability (Kalam et al., 2020; Selim, Abdelhamid, and Mohamed, 2021; Chen et al., 2021). This metabolic diversity likely enabled their persistence and abundance across the different redox conditions, highlighting their adaptability and ecological versatility.

The large presence of cosmopolitan viruses (vOTUs present across all redox conditions) suggests that these viruses may be generalists or infect hosts with significant metabolic plasticity. Since our study utilized metagenomes, which capture a representative subset of the viral community—often reflecting proviruses or viruses infecting the most abundant microbes—this may account for the observation that approximately a third of the vOTUs are found in every redox condition. We examined the virus-host linkages of the cosmopolitan vOTUs and found that nearly all of their predicted hosts belonged to the bacterial phyla *Pseudomonadota*, *Acidobacteriota*, and *Actinomycetota*. The remaining predicted hosts were from Bacillota (formally Firmicutes), Myxococcota, *Planctomycetota*, and *Verrucomicrobiota*, which are also known for their diverse metabolic capabilities and were present across all redox conditions (Chen et al., 2021). This distribution indicates that the cosmopolitan viruses are likely to infect versatile and adaptable hosts, able to thrive in varying environmental conditions due to their broad metabolic capabilities. This supports the idea that these viruses target generalist hosts that can survive and proliferate across different redox environments.

A comparison of AMGs carried by cosmopolitan vOTUs and those selected by redox conditions, provides insight into the potential functionalities imparted by viruses within the microbiome. Focusing on glycoside hydrolases from ^13^C-enriched vOTUs, we see vOTUs carry specific AMGs based on the environmental redox condition. Glycoside hydrolases are enzymes that break down polysaccharides into smaller products, making them a key part of carbohydrate degradation and the carbon cycle. Glycoside hydrolase AMGs involved in the degradation of complex carbohydrates, such as xyloglucans and arabinoxylans commonly found in plant cell walls (Vandermarliere et al., 2009), were more abundant in oxic soils. Aerobic metabolism yields more energy, which supports the function of enzymes like GH 16 and GH 43, which are involved in the initial and intermediate stages of plant biomass degradation (Bardgett et al., 2008; Vuong and Wilson, 2010). Conversely, anoxic conditions showed increased glycoside hydrolases that degrade simpler polysaccharides, such as starch (GH 13), oligosaccharides, and disaccharides into glucose (GH 3), supporting anaerobic fermentation pathways. We see a similar trend in the AMGs carried by ^13^C-enriched vOTUs infecting the top two genera, *Bradyrhizobium* and *Mycobacterium*, present across the redox gradient. Except for ABC transporters, we observe fewer AMGs in active vOTUs under anoxic conditions and more under oxic conditions. The lower metabolic activity in the anoxic soils makes it less favorable for expressing a wide range of metabolic functions. AMGs like ABC transport systems are more abundant under anoxic conditions due to their roles in survival strategies and anaerobic metabolism, such as nutrient acquisition and stress response, and energy and electron transport as previously shown (Hernández and Vives, 2020; Trubl et al., 2021). These findings highlight the potential for viruses to contribute to the functional plasticity of their hosts in different redox environments.

## Conclusion

Our study highlights the significant influence of redox conditions on shaping microbial and viral behavior, as well as virus-host interactions in soils. By applying two different assembly methods to both bulk and density-fractionated DNA for stable isotope probing metagenomes, and incubating soils under three distinct redox regimes, we identified unique viral populations and revealed differences in microbial and viral community composition and functionality across these different redox conditions. The anoxic samples exhibited the most distinct viral communities, with an increased potential for modulating host metabolism. Our work showed that the assembly approach influences virus recovery from metagenomes, with co-assembly revealing greater viral contig and vOTU diversity, effectively detecting rare viruses. The presence of ^13^C-enriched vOTUs under both oxic and anoxic conditions, along with redox-specific AMGs, emphasizes the active roles of viruses in plant biomass degradation and their ability to adapt to changing environmental conditions. Our findings indicate that static environments may support a higher viral richness than fluctuating ones and that viral populations can range from highly specialized to broadly adaptable, shaped by the adaptability of their hosts and environmental conditions. These insights are important for understanding how redox conditions affect virus-host interactions and microbial decomposition processes, which are key factors in carbon cycling. By influencing the decomposition of organic matter and the release of greenhouse gases, these processes have significant implications for climate change.

## Acknowledgments

We thank Jose Liquet-Gonzalez, Mike Allen, Jessica Wollard, Heather Dang, Summer Ahmed, Christina Ramon, Christine O’Connell, Shalini Mabery, Rachel Neurath, Jordan Stark, Omar Gutiérrez del Arroyo, Jean Lodge, Sarah Stankavich, Grizelle Gonzalez, and Sharon Rodríguez for their support in the laboratory, in the field, or with figures, and for providing advice.

## Competing Interests

The authors declare no competing financial interests.

## Ethics approval and consent to participate

Not applicable

## Consent for publication

Not applicable

## Funding

Support for this experiment and data collection was provided by a U.S. Department of Energy Early Career Research Program Award to J. Pett-Ridge (SCW1478) administered by the Office of Biological and Environmental Research (OBER), Genomic Sciences Program. Sequencing was conducted via the Joint Genome Institute Community Sequencing Program, Proposal ID 502924 to J. Pett-Ridge. Site support in Puerto Rico was provided by the Luquillo CZO (EAR-1331841) and LTER (DEB-0620910). Additional support for data analysis was provided by LLNL’s U.S. Department of Energy, Office of Biological and Environmental Research, Genomic Science Program LLNL ‘Microbes Persist’ Scientific Focus Area (#SCW1632) and a Laboratory Directed Research & Development grant (18-ERD-041) to S.J.B. All work at Lawrence Livermore National Laboratory was performed under the auspices of U.S. Department of Energy under Contract DE-AC52-07NA27344. Work conducted by the U.S. Department of Energy Joint Genome Institute (https://ror.org/04xm1d337) (JGI award 10.46936/10.25585/60000880), a DOE Office of Science User Facility, was supported by the Office of Science of the U.S. Department of Energy operated under Contract No. DE-AC02-05CH11231.

## Authors’ contributions

G.T. analyzed the data, prepared figures, and wrote the initial draft of the manuscript. I.L. analyzed the data, prepared figures, and contributed to writing. A.C. collected samples, designed the study, set up the incubations, made sequencing decisions, analyzed microbial sequence data and edited the writing. J.A.K. analyzed SIP microbial sequence data and contributed to writing. A.B. collected samples, helped to design the study and conduct the incubations, and edited the writing. R.R. analyzed microbial data and edited the writing. R.M. analyzed the SIP sequence data and edited the writing. S.J.B acquired funding and oversaw the SIP sample processing and edited the manuscript. J.P-R. acquired funding and supervised the project, conceptualized the study, helped write and edit the manuscript, and served as senior author. All co-authors helped with interpretation of data and approved the final draft submitted.

## Notes

### Competing Interest Statement

The authors have declared no competing interest.

