## Supplementary figures and images for "Soil redox drives virus-host community dynamics and plant biomass degradation in tropical rainforest soils"

### Supplemental Figure 1

A)

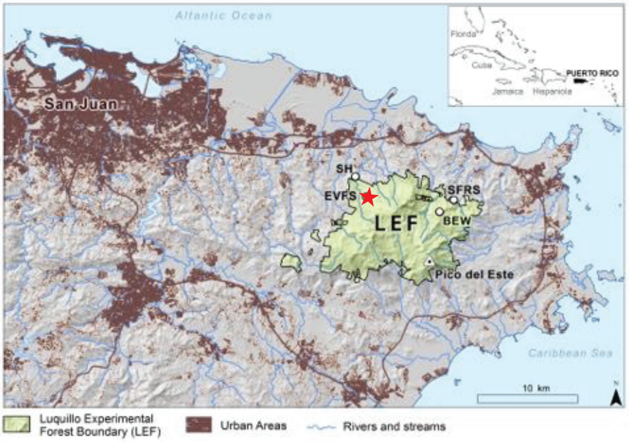


El Verde Field Station (EVFS)

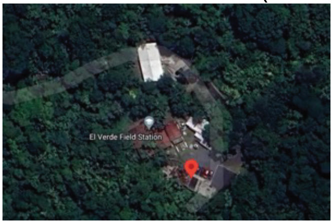


B)

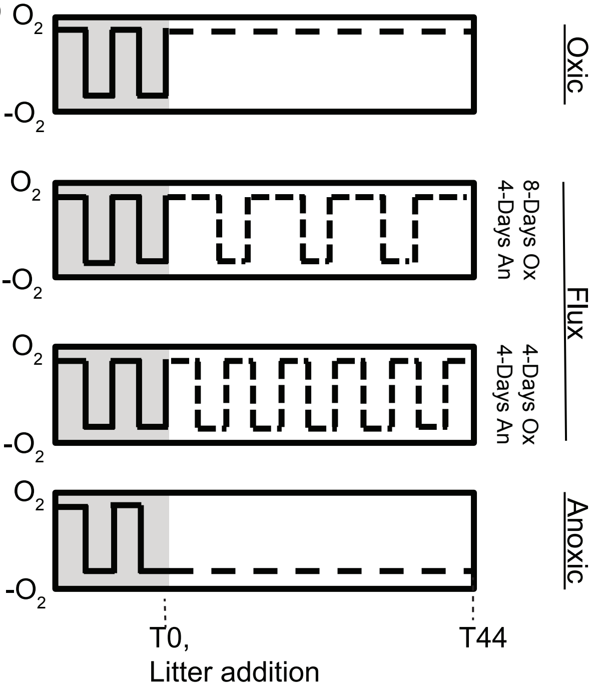

### Supplemental Figure 2

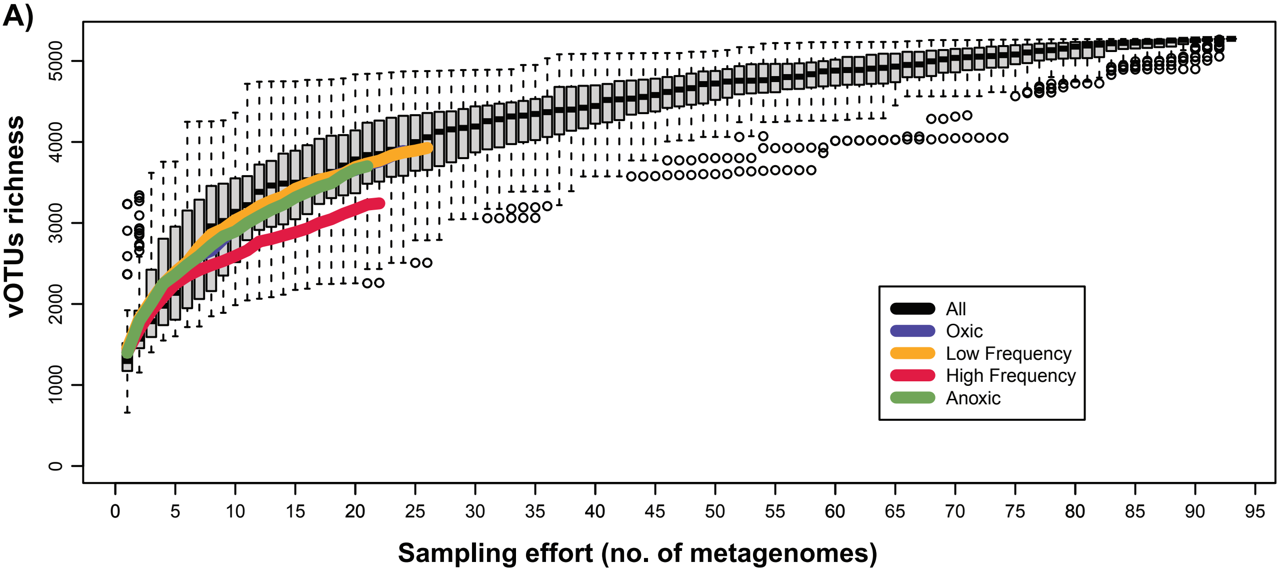

### Supplemental Figure 3

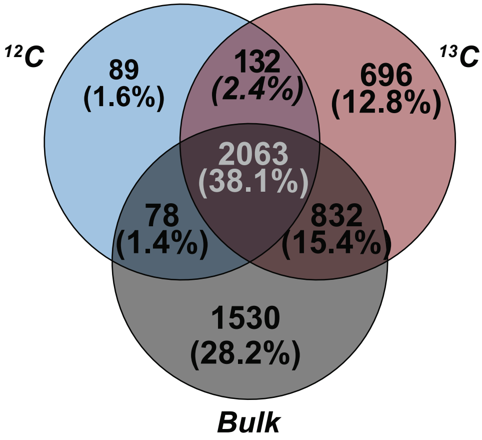

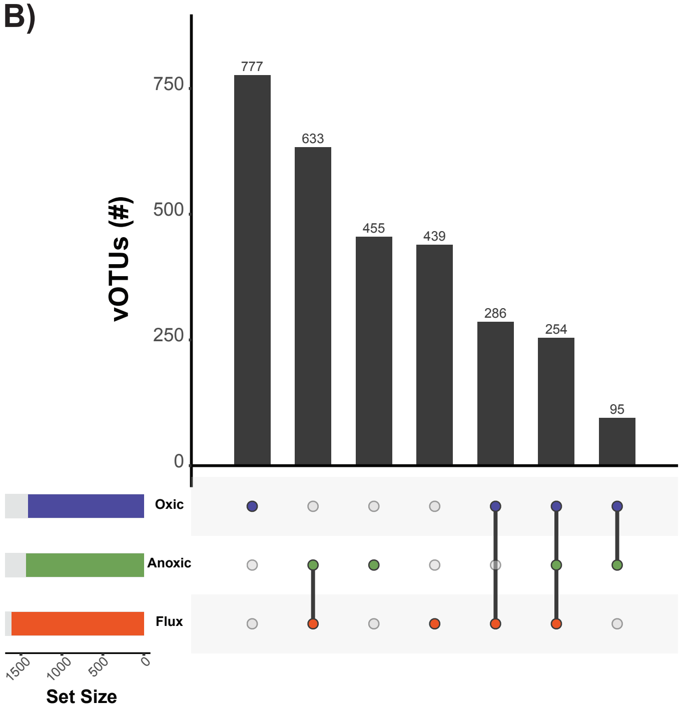

### Supplemental Figure 4

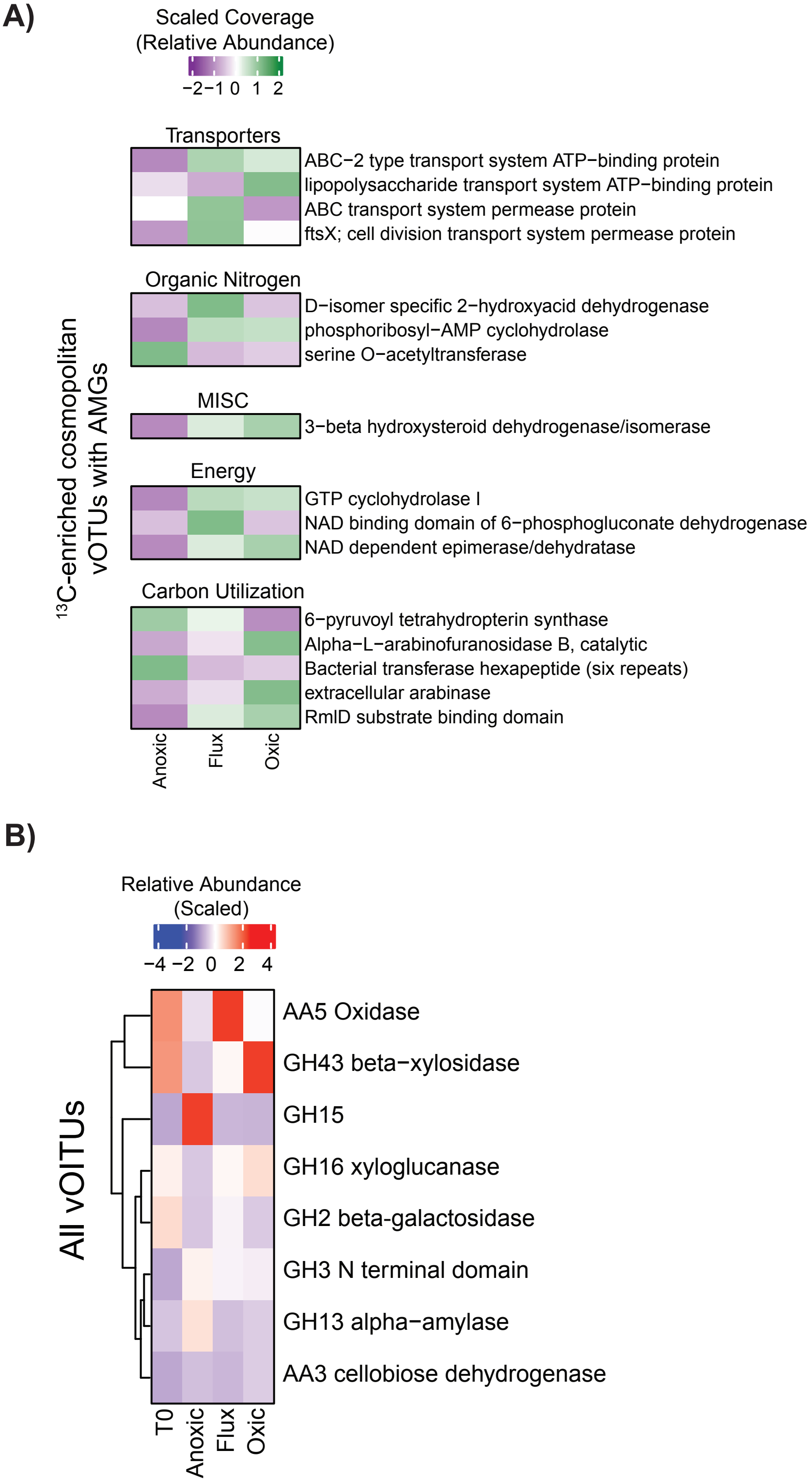
